# Persistent *Staphylococcus aureus* in human musculoskeletal tissues despite targeted antibiotic therapy

**DOI:** 10.64898/2026.06.03.728378

**Authors:** Benedict Morin, Vishwachi Tripathi, Anne-Kathrin Woischnig, Joana Erdmann, Aya Iizuka, Daniel Baumhoer, Katharina Rentsch, Loïc Sauter, Ewelina M. Bartoszek, Peter Keller, Krittapas Jantarug, Pablo Rivera-Fuentes, Baptiste Hamelin, Salome Hosch, Kirsten Mertz, Mario Morgenstern, Martin Clauss, Dirk Bumann, Richard Kuehl, Nina Khanna

## Abstract

*Staphylococcus aureus* is a leading cause of musculoskeletal infection with frequent recurrence despite surgery and antibiotic therapy. Its persistence in human tissues under antibiotic pressure remains unclear. We performed longitudinal *in situ* analyses of matched tissue samples before and during therapy from patients undergoing repeated surgery using advanced imaging and tissue-based metagenomics. *S. aureus* remained detectable in all samples—including culture-negative specimens—and whole-genome sequencing confirmed persistence of the infecting strain. Antibiotic initiation reduced bacterial burden by 1.05log_10_ bacteria/mm^3^ but residual loads stabilized thereafter. Quantitative imaging identified a culture detection threshold (∼10^3^bacteria/mm^3^), indicating that culture negativity reflects diagnostic sensitivity limits rather than true bacterial clearance. Persistence under antibiotic therapy was associated with high baseline burden and intracellular sequestration, whereas long-term persistence was characterized by predominantly extracellular bacteria capable of driving relapse. These findings reveal sustained tissue persistence despite apparent microbiological clearance and guide precision strategies to target resilient bacterial reservoirs.

## Main

*Staphylococcus aureus (S. aureus)* remains a leading cause of infection-related morbidity and mortality.^1–3^ It can infect most human tissues including bone and joints, causing osteomyelitis, septic arthritis, fracture-related infections (FRls), and prosthetic joint infections (PJls). Incidence is rising with population aging and increased implant use.^3,4^ Despite guideline-based surgical and antimicrobial therapy, infection persistence and/or relapse occur in up to 10-50% of cases.^3,5–7^ Laboratory studies propose survival strategies including biofilms, persisters and intracellular evasion.^8–11^ However, their clinical relevance remains unclear due to limited human data and the inability of routine diagnostics and histopathology to detect these phenotypes.^12–18^

To address this gap, we established a longitudinal retrospective cohort of patients undergoing repeated surgeries at the same anatomical site for musculoskeletal. *aureus* infection with tissue sampling before antibiotic initiation and during or after guideline-concordant therapy. Among 2,176 cases of musculoskeletal *S. aureus* infection screened, we identified 14 patients with matched longitudinal tissue samples **(Fig. 1a)**.

**Fig. 1.**
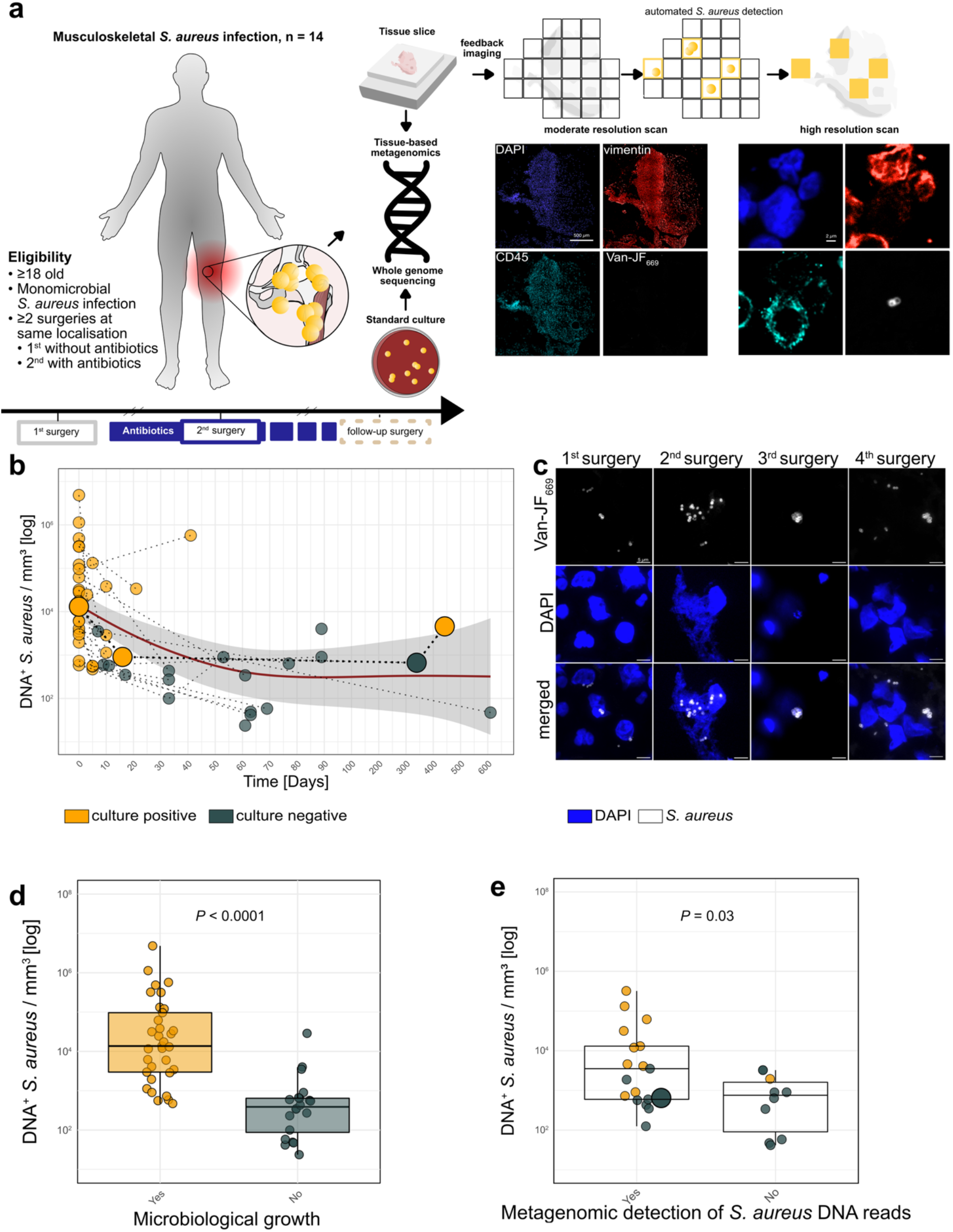
Study design and persistence of *S. aureus*. **a**, Study flow diagram with representative images of moderate and high resolution scans. **b**, DNA^+^ *S. aureus*/mm^3^ across sequential surgeries. Lines connect matched samples from the same patient and anatomical site. Red line, generalized additive model (GAM); grey shading, 95% Cl. The index patient is highlighted in bold. **c**, Representative confocal images of *S. aureus* from sequential surgeries in the index patient. **d**, DNA^+^ *S. aureus*/mm^3^ stratified by culture result. P-value by Wilcoxon rank-sum test (log transformed data). **e**, DNA^+^ *S. aureus*/mm^3^ stratified by metagenomic detection of *S. aureus* reads. P-value by Wilcoxon rank-sum test (log-transformed data). Reads from 3^rd^ surgery of the index patient in bold.

As a representative example, we describe a patient who developed 5. *aureus* septic arthritis of the knee. Initial surgical debridement and intravenous cefazolin were followed by a second debridement two weeks later for persistent infection, again yielding 5. *aureus*. After six weeks of antibiotic therapy, infection resolved clinically and microbiologically. Forty-eight weeks later, total knee arthroplasty was performed to treat post-infectious joint destruction. At this third surgery, *S. aureus* was not suspected; cultures were negative, and the patient recovered. Fifteen weeks later, infection recurred, confirmed by intraoperative isolation of *S. aureus* (4^th^ surgery; **Suppl. Fig. 1a)**. Whole-genome sequencing of cultured isolates from the first, second, and fourth surgeries revealed minimal variation (3-14 single-nucleotide polymorphisms), confirming persistence of the original strain **(Suppl. Fig. 1b)**. These findings suggest that despite adequate therapy and debridement, *S. aureus* survives long term in host tissue, forming a relapse reservoir.

To systematically assess long-term bacterial persistence, we analyzed 54 tissue samples from the 14 patients. Median age at diagnosis was 63 years (IQR 55-69), and 36% were female. In total, 9 patients (64%) underwent two surgeries, 3 (21%) underwent three, and one patient each (7%) required four and five surgeries. Infection types included PJls (57%), FRIs (14%), and native osteomyelitis (14%) **(Suppl. Table 1)**. During the initial surgery, all conventional cultures were positive. In contrast, in sequential samples, standard cultures were negative in 18 samples obtained during 13 surgical procedures, 9 of which were collected while patients were clinically asymptomatic. To overcome limited culture sensitivity, we used high-resolution confocal imaging with the *S. aureus* probe Van-JF_669_ and counterstains for DNA (DAPI), leukocytes (CD45), and cytoskeleton (vimentin), with >95% agreement to manual classification.^19,20^ Across all 54 samples, we identified 22,861 individual *S. aureus* cells, with DNA^+^ bacteria detected in every sample, including all culture-negative specimens. Second surgeries occurred after a median of 10 days (IQR, 6-31), follow-up surgeries 77 days (IQR, 43-214) after the initial procedure **(Fig. 1b,c)**.

Median bacterial density was 1.6×10^4^/mm^3^ (IQR, 5.5×10^3^-1.0×10^5^) before antibiotic therapy and decreased to 5.9×10^2^/mm^3^ (IQR, 4.1×10^2^-1.4×10^4^; *P* = 0.001) after treatment initiation at the second surgery, remaining stable in subsequent procedures *(P* = 0.4) (**Fig. 1b; Suppl. Fig. 1c,d)**. In paired tissue analyses, antibiotic initiation was associated with a median reduction of 1.05 log_10_ (IQR, 0.42-1.22), whereas no further significant decline was observed between the second and follow-up surgeries (median 0.30 log_10_). Median antibiotic exposure was 10 days (IQR 5-29) at the second surgery and 53 days (IQR, 33-66) at follow-up surgeries (**Suppl. Fig. 1d)**.

The proportion of DNA^+^ bacteria decreased from 81% (IQR, 52-91) to 45% (IQR, 28-65; *P* = 0.01) during therapy but returned to 99% (IQR, 57-100; *P* = 0.03) in later surgeries **(Suppl. Fig. 1d)**.

These findings indicate that antibiotic therapy affects bacterial viability in tissues yet fails to sufficiently reduce total bacterial burden despite the isolates being highly susceptible in time kill assays and antibiotic concentrations in infected tissues exceeding the minimum inhibitory concentration by more than 20-fold **(Suppl. Fig. 1e-g)**. Notably, the initial decline likely overestimates antibiotic efficacy, as surgical debridement removes the primary infected tissue reservoir.

Across all patients, amount of DNA^+^ *S. aureus* correlated with culture positivity *(P* <0.0001) **(Fig. 1d)**. Receiver operating characteristic (ROC) analysis established a culture detection threshold of ∼1.1×10^3^ DNA^+^ bacteria/mm^3^ (AUC 0.9; 88% sensitivity, 86% specificity), showing that subculture-level persistence is common, and absence of culture growth reflected diagnostic detection limits rather than microbiological eradication.

We performed tissue-based metagenomic sequencing as an orthogonal approach to validate the presence of locally persisting bacteria as a potential reservoir for relapse. In the index patient, metagenomics confirmed culture-silent persistence during the clinically quiescent phase, detecting 38 5. *aureus* reads in culture-negative tissue. When extended across additional tissue samples, metagenomic read counts correlated with the burden of DNA^+^ *S. aureus (P* = 0.03, **Fig. 1e)**, reinforcing that subculture-level persistence is both detectable and biologically relevant.^21^

Antibiotic therapy reduced the number of DNA^+^ bacteria but was not associated with a consistent predominance of specific morphological patterns, such as large aggregates, bacteria associated with extracellular DNA, or predominantly intracellular remnants. Histological appearances remained heterogeneous and patient-specific, without enrichment of distinct morphologies **(Suppl. Fig. 1h-i**, confirming our previous observations^20^).

We next examined whether bacterial behavior under antibiotic therapy differed between patients with microbiological failure (growth persistence) and those achieving clearance at the time of second surgery. *S. aureus* remained culture-positive in 8 of 14 patients despite initial surgical intervention and antibiotic treatment **(Fig. 2a)**. The persistence and clearance groups were comparable with respect to age, sex, comorbidity burden, antibiotic treatment duration, and the interval between first and second surgeries. The principal clinical distinction was that all cases with growth clearance occurred in PJI, possibly reflecting a lower threshold for reoperation **(Suppl. Table 2)**.

**Fig. 2.**
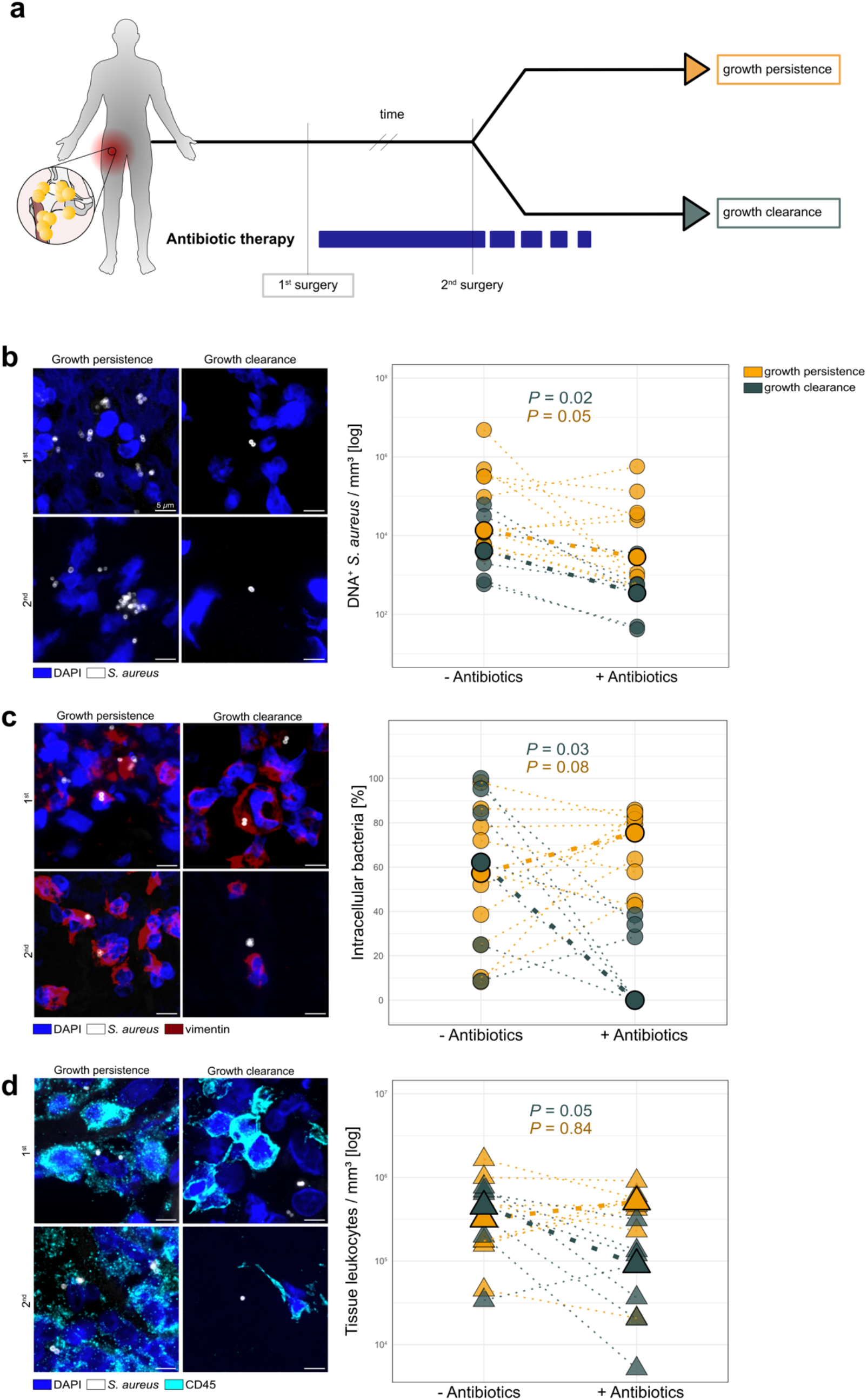
Intracellular *S. aureus* drives growth persistence under antibiotic therapy. **a**, Schematic of study comparison between growth persistence and clearance at 2^nd^ surgery. **b**, Left panel: Representative tissue images from one patient with persistence and one with clearance, and quantification of DNA^+^ 5. *aureus/*mm^3^ at 1st and 2nd surgeries. Right panel: Lines connect matched longitudinal samples. Medians indicated. P-values by Wilcoxon rank sum test (log-transformed data). **c**, Left panel: Representative images of intracellular and extracellular *S. aureus*. Right panel: Proportion of intracellular bacteria by antibiotics. P-values by Wilcoxon signed-rank test. **d**, Left panel: Representative image of S. *aureus* within immune cell-rich regions. Right panel: Leukocyte density by surgery and antibiotic status. P-values by Wilcoxon rank-sum test (log-transformed data).

Baseline tissue *S. aureus* burden tended to be higher in patients with growth persistence than in those achieving clearance (median 1.4×10^5^ [IQR, 8.9×10^3^-3.2×10^5^) vs. 4.1×10^3^ [1.3×10^3^−2.2×10^4^] bacteria/mm3; *P* = 0.06; **Fig. 2b; Suppl. Table 2)**. In the persistence group, antibiotic therapy led to heterogeneous responses and overall reduction in bacterial density to 2.8×10^3^/mm^3^ (IQR, 7.3×10^2^-3.6×10^4^; *P* = 0.05). In contrast, the clearance group exhibited a decrease across all patients, from 4.1×10^3^ to 3.5×10^2^ bacteria/mm^3^ (IQR, 2.0×10^2^-5.8×10^2^; *P* = 0.02) **(Fig. 2b)**. Although relative reductions were similar between groups, persistence occurred when high bacterial burdens remained above the threshold required for culture negativity after therapy, whereas lower baseline loads allowed comparable proportional reductions to result in microbiological clearance.

Intracellular localization of *S. aureus* increased in cases with growth persistence, from 57% (IQR, 32-75) to 76% (IQR, 60-82; *P* = 0.08), whereas it declined in cases achieving clearance, from 62% (IQR, 42-90) to 0% (IQR, 0-31; *P* = 0.03) **(Fig. 2c)**. Immune cell densities remained stable in the persistence group between the first and second surgeries (median 3.2×10^5^ [IQR, 1.7×10^5^-6.1×10^5^)vs. 5.1×10^5^ [3.7×10^5^-5.7×10^5^)CD45^+^ cells/mm^3^; *P* = 0.8), but declined in the clearance group from 4.5×10^5^ (IQR, 2.7×10^5^-6.7×10^5^) to 9.1×10^4^ cells/mm^3^ (IQR, 2.8×10^4^-1.3×10^5^; *P* = 0.05) **(Fig. 2d)**. These findings indicate that growth persistence under antibiotic therapy is associated with sustained intracellular sequestration and a lack of immune cell mediated reduction, whereas successful clearance coincides with loss of intracellular bacteria and resolution of the inflammatory infiltrate. In contrast, during long-term persistence, extracellular bacteria may predominate and sustain infection relapse, as illustrated by the index patient **(Suppl. Fig. 2)**.

This novel longitudinal human study provides direct *in situ* evidence that *S. aureus* persists in musculoskeletal tissue for extended periods despite targeted antibiotic therapy. The observation that bacterial burden increased under therapy in only one patient indicates that antibiotic treatment exerts effective growth suppression *in vivo*. However, this effect was insufficient for bacterial eradication, resulting in only modest reduction in bacterial load between the first and second surgeries and no additional decrease thereafter. Importantly, this initial reduction is likely overestimated, as surgical debridement removes heavily infected tissue, leaving subsequent biopsies to sample predominantly residual, less densely colonized tissue. Collectively, these findings demonstrate that antibiotic therapy alone fails to eradicate *S. aureus* from infected musculoskeletal tissue.

Quantitative imaging detected residual bacteria in all samples, including culture-negative tissue, highlighting the need for more sensitive diagnostics capturing low-level persistence and indicating that negative cultures primarily reflect methodological detection limits rather than true bacterial clearance. The identified threshold of ∼10^3^ DNA^+^ bacteria/mm^3^ provides a quantitative reference for evaluating new diagnostics and contextualizing microbial burden in relation to clinical relevance and local host response.

In cases of bacterial growth persistence under antibiotic therapy, our data imply that high initial tissue burden and intracellular bacterial sequestration rather than resistance selection or large cluster formation drive early persistence. Standard antimicrobial regimens for *5. aureus* infections and the agents predominantly used in our cohort, are known to have limited intracellular activity, whereas antibiotics with enhanced intracellular efficacy may offer therapeutic advantages that warrant clinical validation.^22−25^ Durable eradication will likely require strategies combining intracellularly active antimicrobials with host-directed approaches designed to enhance phagocytic clearance or disrupt protective tissue reservoirs. Importantly, while intracellular bacteria appear to sustain early persistence under antibiotic pressure, extracellular bacteria predominate during long-term persistence and may serve as a source of relapse.^26,27^ Beyond bacterial burden and localization, additional host and pathogen determinants are likely to modulate persistence and merit systematic investigation.

While the longitudinal *in situ* sampling constitutes a major strength of this study, the single center design and small, highly selected cohort of 14 patients may limit generalizability and require validation in larger prospective cohorts. In addition, interpreting DNA^+^ *S. aureus* as viable bacteria may be a limitation. Although DNA detection alone does not prove viability, the sustained presence of membrane-intact bacteria, lack of degradation over time, and association with relapse support the presence of viable organisms.^28,29^

Together, these findings redefine the pathobiological framework of *S. aureus* musculoskeletal infection by linking tissue-level bacterial burden and spatial compartmentalization to therapeutic failure and relapse, thereby providing a rational foundation for precision strategies targeting resilient bacterial reservoirs.

## Methods

### Patient sampling

Tissue samples were collected during surgery by orthopaedic surgeons specialized in musculoskeletal infections between 2013 and 2024 at the University Hospital of Basel, Switzerland. Most of the surgeries were performed by the same two surgeons (M. C. and M. M.) both specialized in musculoskeletal infections. Samples consisted either of infected soft tissue, bone or abscess.

For every tissue localization, one sample was split into two. One for microbiological diagnostics and one for pathology. In microbiological diagnostics, the samples were streaked on Mueller Hinton Agar plates and put into liquid medium for the purpose of bacterial culture and streaked on a microscopy slide and stained with gram for microscopic detection. In the pathology, samples were fixed in 4% paraformaldehyde, embedded in paraffin and either directly visualized or stored.

The study was approved by the local Ethical Review Board (Ethikkommission Nordwestschweiz, Project-ID 2020-02588) and performed in compliance with ethical regulations.

### Patient screening for imaging

35’143 patients who have received an infectious disease consultation within the last 10 years were screened. 2’176 patients had secured or probable monomicrobial infection with *S. aureus*. Out of those, we excluded patients with immunosuppressive disease and/or therapy and finally included 14 patients with monomicrobial musculoskeletal *S. aureus* infection receiving ≥ 2 consecutive surgeries at the same site: one without antibiotic treatment 2 ≥ weeks before the surgery and one with prior antibiotic treatment.

### Measurement of antibiotic concentration in interstitial fluid

Unbound interstitial fluid concentrations in tissue samples were determined using equilibrium dialysis. 15 mg tissue sample was added on top of a membrane with 10k molecular weight cut-off (Slide-A-Lyzer™ MINI Dialysis Devices, Thermo Fisher Scientific, Reinach, Switzerland) and approximately 18 µL of a macromolecular isotonic solution (Voluven®, Fresenius Kabi, Kriens, Switzerland) was placed under the membrane. The device was put in a 1.5 mL microcentrifuge tube and incubated at 37 °C for 5 hours. Following the incubation and sample preparation steps, which included homogenization and protein precipitation, flucloxacillin concentrations were determined in the residue and the dialysate. The analysis was performed using high-performance liquid chromatography coupled to tandem mass spectrometry. The free tissue fraction was calculated from the ratio of the measured dialysate and residue concentrations. The unbound interstitial fluid concentration was obtained by multiplying the free tissue fraction by the total tissue concentration which was assessed in a second aliquot of 10 mg tissue.^30^

### Tissue staining

4 µm formalin-fixed paraffin-embedded (FFPE) musculoskeletal tissue samples were deparaffinized and rehydrated, including two 10-min washes in 100% xylene at room temperature (RT), followed by serial incubations at RT: two 5-min incubations in 100% ethanol, and 2 min each in 96%, 70%, and 50% ethanol, ending with distilled water. For antigen retrieval, tissues were incubated in EDTA-based epitope retrieval buffer (pH 9.0, AR9640, Biosystems) for 20 min at 95°C, then cooled for 15 min at RT. Slides were transferred to PBST (PBS+ 0.1% Tween 20 (Merck)) for permeabilization.

Tissue structure was circled with a hydrophobic pen (ab2601, Abcam), and 250 µL blocking buffer (PBS+ 0.1% Tween 20 + 5% goat serum (ab7481, Abcam)) was added, then incubated for 1 hour at RT in a dark humidity chamber. Blocking buffer was removed by tilting, and primary antibodies in blocking buffer were applied overnight at 4°C in the dark. Primary antibodies included rabbit anti-human CD45 (1:100, ab30470, Abcam) and mouse anti-human Vimentin (1:200, ab89789, Abcam).

The following morning, primary antibodies were removed by tilting and rinsing with PBST (3x), followed by three 5-min washes in PBST. Secondary antibodies were added (goat anti-rabbit Alexa Fluor (AF) 555, A-21422, and goat anti-mouse Alexa Fluor (AF) 750, A-21037, Thermo Fisher Scientific, both 1:500 in blocking buffer) and incubated for 1 hour at RT in the dark.

Secondary antibodies were removed with PBST (3x), followed by three 5-min washes in PBS. For bacterial staining, Van-JF_669_ ^19^ (1.66 µg/mL) and DAPI (1.25 µg/mL, Abcam, ab228549) in PBS were applied and incubated for 15 min at RT in the dark. Slides were rinsed (3x) and washed in PBS (3x 5 min) at RT in the dark. Finally, slides were mounted with two drops of aqueous mounting medium (F4680, Merck) and covered with #1.5 glass coverslips (Novoglas).

### Image acquisition

Fluorescence labels used were DAPI (Abcam, ab228549), AF555 (Abcam, ab150078), Janelia Fluor 669 (JF_669_)^19^ and AF750 (Thermo Fisher Scientific, A-21037). Images were collected with a Nikon Ti2-E inverted microscope, equipped with a Crest V3 spinning disk unit (CrestOptics, pinhole size = 50 µm), using a Plan Apo Lambda S 40x Silicone (NA = 1.25), and a Plan Apo Lambda S 100x Silicone (NA= 1.35) objective. DAPI, tissue autofluorescence, AF555, JF_669_ and AF750 fluorescence were excited with a Celesta (Lumencor) at 405, 510, 546, 640 and 749 nm, respectively, and collected with a penta-edge 421/491/567/659/776 dichroic beam splitter, a dual-edge 471/539 splitter, and 438/24 (DAPI), 511/20 (autofluorescence), 560/25 (AF555), 685/40 (JF_669_) single-bandpass filters and a 441/30, 511/26, 593/36, 684/34, 817/66 (AF750) penta-bandpass filter. Images were acquired with a Photometrics Kinetix camera controlled with the Nikon NIS software. Excitation light intensity and camera exposure were adjusted for each channel to prevent image oversaturation and were kept the same for samples from the same patients.

The NIS JOBS module was used for automated image acquisition. Briefly, a tiled overview image (15% tile-overlap) was captured using the 40x objective for all the channels for a total of 9 z-planes. The Nikon Perfect Focus System was used to keep the focus over large areas, and Z-stacks were acquired in 0.5 µm step size. Subsequently, maximum intensity projections of the tiles were stitched for identification of bacterial objects. The NIS GA3 module was used for object identification, which was performed as follows: the gaussian blur (sigma = 1) of the autofluorescence channel was subtracted from the JF_669_ channel to obtain a background corrected image of the bacteria. Then a threshold manually adjusted per image in the range from 3 - 10 for the JF669 channel defined objects of interest. Tiles with objects of interest in its center were rescanned using the 100X objective. For that, auto-focusing (PFS) on the JF_669_ was performed, and Z-stacks were acquired over 13 planes in step size 0.3 µm around the focus plane for each channel.

### Image analysis

Analysis of stitched moderate resolution 40X scans was performed on QuPath, Version 0.5.1.^31^ The tissue was manually outlined and cell detection was performed using the Watershed Cell Detection plugin on the DAPI channel, using pixel size = 0.5, background radius = 2, background by reconstruction = true, median radius = 0, sigma = 1.5, min area = µm^2,^ max area = 500 µm^2,^ threshold = 1, split by shape = false, cell expansion = 5 µm, include nuclei = true, smooth boundaries = true, and make measurements = true. Cells were then classified as CD45^+^ according to the mean intensity of the autofluorescence (YFP) channel within the cell compartment. The mean intensity threshold was adjusted per tissue sample but kept the same for tissues originating from the same patient. Bacteria were segmented using an OpenCV-based pixel classifier trained on the JF_669_ channel. The intensity threshold for classifying pixels as bacteria versus background was adjusted for each tissue sample to account for variation in signal and kept consistent for samples from the same patient.

Eventually, the cell detection measurements were exported, as well as the annotation measurements. The annotation measurements were used to extract the number of CD45^+/−^ cells per tissue region. The entries for bacterial annotations were manually extended, by visually inspecting and validating each bacterial object (i.e. classified as true positive/negative) and approximate count of number of bacteria per object. In addition, each bacterial object was visually classified in respect to its niche (intracellular / extracellular), within CD45^+/−^, or surrounding environment (not within a cell object).

Finally, counts of intracellular, extracellular, and tissue-residing bacteria and the number of CD45+ cells per tissue volume were exported.

Analysis of 100X high-resolution 3D images were performed in (Fiji is Just) lmageJ, 2.14.0^32,^ NIS-Elements AR Analysis 5.42.06 (Nikon), or directly on the data storage platform OMERO (hosted by the IMCF, Biozentrum University of Basel). In tissues with high bacterial abundance, at least 100 bacteria per tissue were classified, in tissues with low abundance, all were classified: Briefly, every Van-JF_669_^+^ detection in the center of the image was visually confirmed as bacteria. Then, the number of bacteria per cluster and its niche were described. For the definition of DNA positivity, JF669 and DAPI (inverted grayscale) were simultaneously checked. Included bacteria were noted with a region of interest in OMERO.

The data was further stored in Microsoft Excel, Version 16.89.1 and analysed in Rstudio, Version 2024.04.2+764.

### Genome sequencing and assembly

All three isolates were sequenced with Illumina, and the first isolate was additionally sequenced with Nanopore to generate a complete reference. Raw reads were filtered using Filtlong (v0.2.1) and trimmed using Porechop (v 0.2.4). Genome assembly of the reference was performed using the Flye assembler3^3^ and was polished with RACON^34^ (v1.5.0) and Medaka (v2.2.0). The assembly was further polished with Illumina short-reads using Pilon^35^ (1.23). Completeness was assessed using BUSCO (https://busco.ezlab.org) and correctness using QUAST^36.^

### Metagenomic sequencing

Formalin fixed paraffin embedded tissue was processed for DNA extraction on the EZ1/2 DNA Tissue Kit (Qiagen, 953034) on the EZ2 Connect (Qiagen, 9003210) automate. DNA concentrations were measured using the Qubit dsDNA HS Assay (Thermo Fisher Scientific, 032854) on a DeNovix QFX fluorometer. Library preparation was performed with the Ion Xpress Plus Fragment Library kit (Thermo Fisher Scientific, 4471269) and sequenced on a Thermo Fisher Scientific Ion Torrent S5XL (10M reads per library). Bioinformatics analysis and taxonomic profiling were performed with the Qiagen CLC Genomics Workbench (25.0.1).^21^

### Time-kill curve

To characterize the time-dependent activity of cefazolin and flucloxacillin against planktonic S. *aureus*, we established a semi-automated robotic time-kill assay using the OT-2 liquid-handling platform (Opentrons, USA). To approximate bacterial burdens typically observed in patients with S. *aureus* infections, bacterial suspensions were adjusted to 2 x 10^3^ CFU/ml. Clinical S. *aureus* isolates retrieved from 6/14 patients were exposed to either cefazolin or flucloxacillin at 10 µg/mL (Cmax).^37^ Parallel growth controls without antibiotic exposure were included to quantify baseline proliferation.

At 0, 2, 4, 6, and 24 hours, the OT-2 robot performed serial dilutions of culture aliquots and automated plating onto Onewell plates (Greiner, Germany) containing Mueller-Hinton agar (MHA; BD, Switzerland). Plates were incubated overnight at 37 °C. Imaging was performed using a FUSION FX7 EDGE system (Vilber Lourmat, France), and CFU counts were obtained using an in-house automated pipeline built in ImageJ/Fiji (v2.9.0).

### SNP analysis and phylogenetic tree

The genetic relatedness of S. *aureus* isolates was assessed by single-nucleotide polymorphism (SNP) analysis. Raw Illumina reads were aligned to the complete reference genome assembled from the first isolate using snippy (v4.6.0). A core genome alignment was generated from the Snippy outputs using snippy-core, retaining only positions present across all isolates. The resulting core SNP alignment was used as input for phylogenetic inference in IQ-TREE2 (v2.4.0). S. *aureus* from a different clonal complex were added to add genetic context and demonstrate that the three patient isolates from this study are monophyletic.

### Statistical analysis

Data are reported as median ± IQR. Statistical significance was determined using the Wilkoxon rank-sum test, or Wilcoxon signed-rank Test, on original or log-transformed data, as indicated. Differences were considered statistically significant if *P* < 0.05. Correlations were evaluated using Spearman rank-order correlation. Dependencies are displayed with a generalized additive model (GAM). Optimal thresholds of the ROC curve were calculated using the Youden Index, unless stated otherwise. All computational analysis and figures were performed in RStudio, v. 2024.04.2+764.

### Declaration of generative Al and Al-assisted technologies in the writing process

Language editing assistance and R code corrections was provided using ChatGPT (OpenAI, San Francisco, CA, USA) for grammar and stylistic refinement. The authors reviewed and edited all AI-assisted outputs and take full responsibility for the content.

---

**Supplementary Fig. 1.**
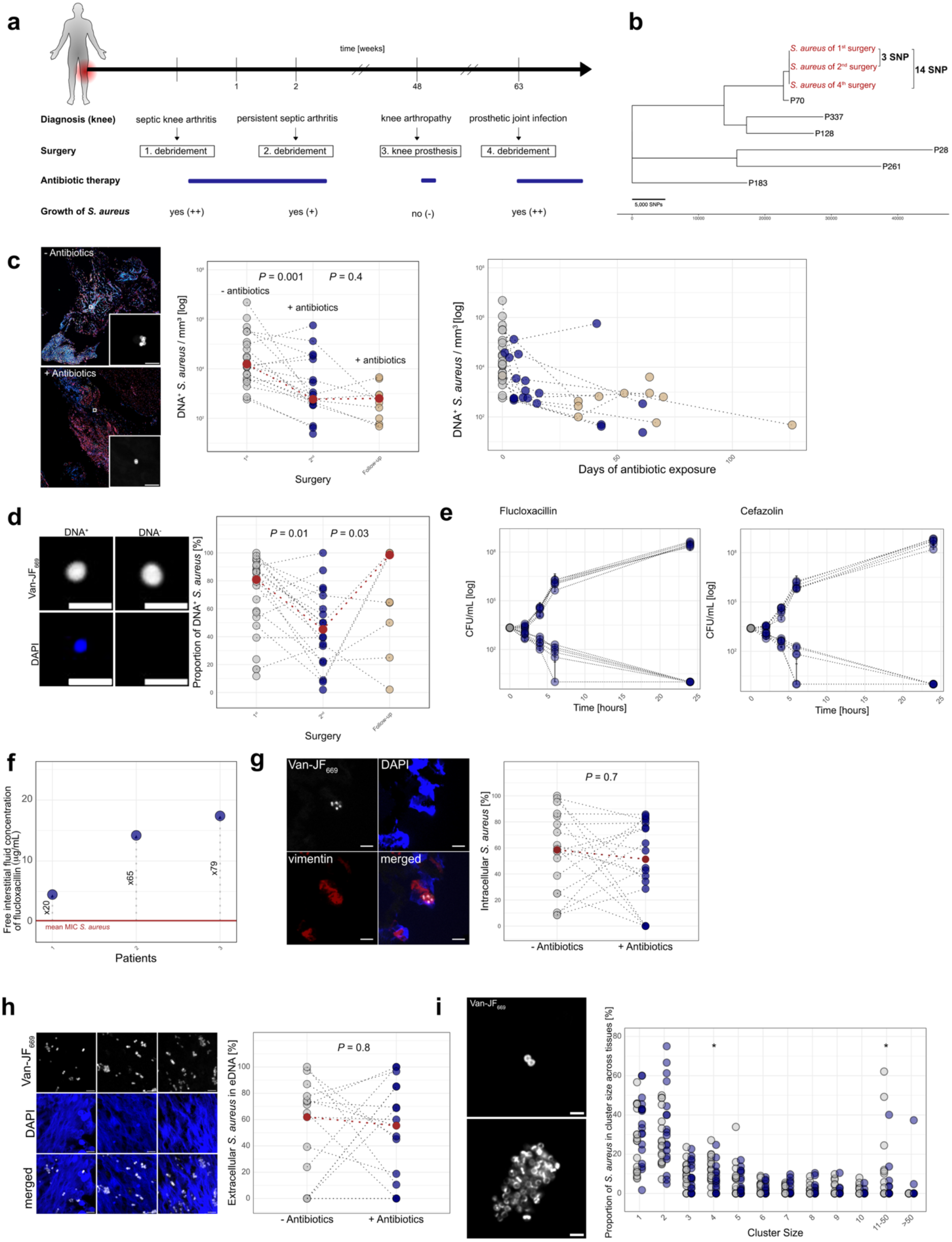
Overview of clinical course, genomic relatedness, imaging, and quantitative analyses of *S. aureus* during antibiotic therapy. **a**, Schematic overview of the patient’s clinical history and timing of surgeries and antibiotic therapy. **b**, Whole-genome phylogenetic tree of *S. aureus* isolates obtained during the first, second, and fourth surgeries, compared with *S. aureus* isolates from other patients with musculoskeletal infections. **c**, Representative images of tissue before and during antibiotic therapy. Scale bar= Sµm; *S. aureus*, grey; DAPI, blue; vimentin, red; CD45, cyan. Quantification of DNA^+^ 5. *aureus*/mm^3^ during first (-antibiotics, grey), second(+ antibiotics, blue) and follow-up surgeries(+ antibiotics, beige) and quantification of DNN *S. aureus/*mm^3^ over antibiotic treatment duration. Each dot represents one tissue sample. Dots are connected when derived from the same patient and tissue type. Red dots indicate median values. P-values were calculated using a Wilcoxon signed-rank test on log-transformed data. **d**, Representative images of DNA^+^ and DNA^−^ *S. aureus*. Scale bar= 2µm; *S. aureus*, grey; DAPI, blue. Proportion of DNA^+^ *S. aureus* across surgeries. Each dot represents one tissue sample. Dots are connected when derived from the same patient and tissue type. Red dots indicate median values. P-values were calculated using a Wilcoxon signed-rank test. **e**, Time-kill curves for six *S. aureus* isolates from the cohort exposed to flucloxacillin or cefazolin. Grey, before antibiotic exposure; blue, under antibiotic therapy. **f**, Free interstitial fluid concentrations of flucloxacillin in three patients with musculoskeletal infections. Each dot represents a single measurement. The red horizontal line indicates the mean minimum inhibitory concentration (MIC) of 5. *aureus* isolates from seven patients. Numbers denote fold-change of tissue concentrations relative to the MIC. **g**, Representative images showing intracellular *S. aureus*. Scale bar= Sµm; *S. aureus*, grey; DAPI, blue; vimentin, red. Proportion of intracellular *S. aureus* by antibiotic status. Each dot represents one tissue sample. Grey dots are tissues from first surgery (- antibiotics), blue dots from second surgery(+ antibiotics). Dots are connected only when derived from the same patient and tissue type. Red dots indicate median values. *P-* value was calculated using a Wilcoxon signed-rank test. **h**, Representative images showing *S. aureus* within extracellular DNA. Scale bar= Sµm. *S. aureus*, grey; DAPI, blue. Proportion of extracellular DNA-associated *S. aureus* by antibiotic status. Grey dots are tissues from first surgery (- antibiotics), blue dots from second surgery (+ antibiotics). Dots are connected only when derived from the same patient and tissue type. Red dots indicate median values. P-value was calculated using a Wilcoxon signed-rank test. **i**, Representative images of large and small *S. aureus* clusters. Scale bar= 2µm. Proportion of 5. *aureus* in different cluster sizes (=number of *S. aureus* per cluster) prior to (grey) or after (blue) initiation of antibiotic therapy. Each dot represents proportions within one tissue sample. Statistical significance of difference in cluster sizes between groups *(P* < 0.05) was evaluated using a Wilcoxon rank-sum test.

**Supplementary Fig. 2.**
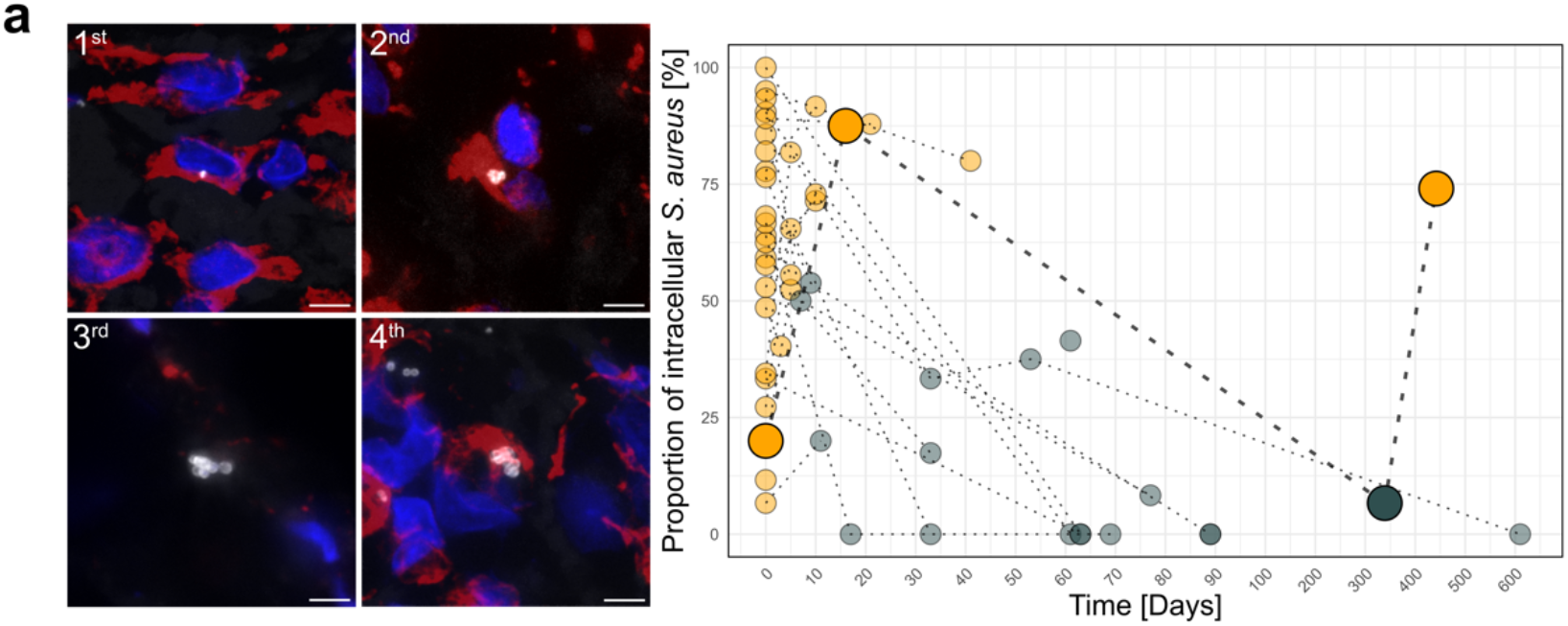
*S. aureus* niches over time in index patient. **a**, Representative images of *S. aureus* niches from tissues during different surgeries. Scale bar= 5µm; *S. aureus*, grey; DAPI, blue; vimentin, red. Proportion of intracellular *S. aureus* over time. Orange points represent tissues with positive culture growth; grey points represent tissues without culture growth. Points are connected only when derived from the same patient, tissue type, and anatomical location. Tissues from the index patient are highlighted in bold.

**Supplementary Table 1.** Patient characteristics at 1^st^ surgery.

| Characteristics | n = 14 |
| --- | --- |
| Age, years median (IQR) | 63 (55-69) |
| Female, n (%) | 5 (36) |
| Diagnosis, n (%) |  |
| Prosthetic joint infection | 8 (57) |
| Fracture related infection | 2 (14) |
| Osteomyelitis without implant | 2 (14) |
| Native joint arthritis | 1 (7) |
| Soft tissue infection | 1 (7) |
| Infection type, n (%) |  |
| Acute | 7 (50) |
| Chronic | 7 (50) |
| Charlson comorbidity index, median (IQR) | 2 (1-3) |

**Supplementary Table 2.** Characteristics of the persistence and clearance group.

| Characteristics | Persistence (n=8) | Clearance (n=6) |
| --- | --- | --- |
| Age, years (IQR) | 59 (50-65) | 66 (62-69) |
| Female, n (%) | 3 (38) | 2 (33) |
| Diagnosis, n (%) |  |  |
| Prosthetic joint infection | 2 (25) | 6 (100) |
| Fracture related infection | 2 (25) |  |
| Osteomyelitis without implant | 2 (25) |  |
| Native joint arthritis | 1 (13) |  |
| Soft tissue infection | 1 (13) |  |
| Infection type, n (%) |  |  |
| Acute | 4 (50) | 3 (50) |
| Chronic | 4 (50) | 3 (50) |
| Charlson comorbidity index, median (IQR) | 2 (1-3) | 3 (1-4) |
| Days until second surgery (IQR) | 10 (5-17) | 14 (10-50) |
| Antibiotic treatment duration prior to 2<sup>nd</sup> surgery (IQR) | 6 (5-12) | 13 (10-36) |
| CRP, mg/dL (IQR) | 5 (3-106) | 80 (39-130) |
| Peripheral leukocytes, 10<sup>9</sup>/L (IQR) | 7 (6-11) | 11 (10-13) |
| Tissue leukocytes/mm<sup>3</sup>(IQR) | 2.8e5 (1.6e5-4.7e5) | 4.5e5 (2.7e5-6.7e5) |
| DNA<sup>+</sup> *S. aureus*/mm<sup>3</sup> (IQR) | 1.4e5 (8.9e3-3.2e5) | 4.1e3 (1.3e3-2.2e4) |
| Proportion of DNA<sup>+</sup> *S. aureus*, % (IQR) | 86 (69-95) | 77 (60-87) |

## Author contributions

B.M., A.K.W., J.E., B.H. and S.H performed experiments; B.M., A.K.W., J.E., A.I., R.K. and N.K. analyzed the data; B.M., V.T., L.S. and E.B. developed the feedback imaging- and analysis workflow; B.M., R. K., D.B., M.C., K.D.M., M.M. and N.K. developed the sampling workflow; B.M., R.K. and N.K. wrote the paper with inputs from all authors.

All data generated or analyzed during this study are included in the manuscript or available upon request.

The authors declare no competing interests.

Correspondence and requests for materials should be addressed to

## Acknowledgements

We thank the DBM microscopy facility for the access to microscopes and help to establish the workflow. We thank the pathology department of the University Hospital of Basel especially Petra Huber, Petra Hirschmann and Daniel Baumhoer for the access to the FFPE slides. We thank the IMCF for providing the OMERO infrastructure and the University of Basel for providing HPC resources (sciCORE) for data storage. We thank Kerstin Strenger, Emre Demirbilek, Anna Sophia Beetschen, Darya Palianina, Dennis Strobbe, Carla Walti, Stefanie Heller, Fanny Linder-Hengy, Claudia Stühler, Sara Zurbrügg, Minia Antelo Varela, Aline Amani Loriol, Jasmin Künnecke, Florian Marro and Noemi Henselmann for their valuable feedback during meetings. Most importantly, we thank the patients allowing us to use their patient material.

